# Enhancing the efficacy of neonicotinoids against mosquitoes and overcoming resistance issues

**DOI:** 10.1101/2023.04.18.537427

**Authors:** Fred A. Ashu, Caroline Fouet, Marilene M. Ambadiang, Véronique Penlap-Beng, Colince Kamdem

## Abstract

**Background:** Neonicotinoids are potential alternatives for targeting pyrethroid-resistant mosquitoes, but their efficacy against malaria vector populations of Sub-Saharan Africa has yet to be investigated. Here we tested and compared the efficacy of four neonicotinoids alone or in combination with a synergist against two major vectors of *Plasmodium*.

**Results:** Using standard bioassays, we first assessed the lethal toxicity of three active ingredients against adults of two susceptible *Anopheles* strains and we determined discriminating doses for monitoring susceptibility in wild populations. We then tested the susceptibility of 5532 *Anopheles* mosquitoes collected from urban and rural areas of Yaoundé, Cameroon, to discriminating doses of acetamiprid, imidacloprid, clothianidin and thiamethoxam. We found that in comparison with some public health insecticides, neonicotinoids have high lethal concentration, LC_99_, reflecting their low toxicity to *Anopheles* mosquitoes. In addition to this reduced toxicity, resistance to the four neonicotinoids tested was detected in *An. gambiae* populations collected from agricultural areas where larvae are intensively exposed to crop-protection neonicotinoids. However, adults of another major vector that occurred in urbanized settings, *An. coluzzii*, were fully susceptible to neonicotinoids except acetamiprid for which 80% mortality was obtained within 72 h of insecticide exposure. Importantly, the cytochrome inhibitor, piperonyl butoxide (PBO), was very effective in enhancing the activity of clothianidin and acetamiprid providing opportunities to create potent neonicotinoid formulations against *Anopheles*.

**Conclusion:** These findings suggest that to successfully repurpose agricultural neonicotinoids for malaria vector control, it is essential to use formulations containing synergists such as PBO or surfactants to ensure optimal efficacy.

## 1. Introduction

The scale up of vector control has been instrumental to the reduction of malaria burden over the last two decades in Sub-Saharan Africa ^1^. Long-lasting insecticidal nets and indoor residual spraying constitute the core vector control interventions and rely on the use of chemical insecticides from 6 classes: pyrethroids, carbamates, organophosphates, organochlorines, neonicotinoids and pyrroles ^2^. Prior to the recent approval of a neonicotinoid and a pyrrole by the World Health Organization (WHO), neurotoxic insecticides that disrupt a sodium channel or inhibit acetylcholinesterase in the insect’s nervous system were widely applied. The similarity of modes of action combined with intensive use of a limited number of active ingredients has created ideal conditions for the emergence and spread of resistance ^3,4^. Indeed, insecticide resistance in malaria vector species has been reported against all the classes of neurotoxic insecticides, posing a challenge to the sustainability of vector control interventions ^5,6^. As a result, the search for new insecticides has become an urgent necessity ^7,8^. In the quest for new active ingredients, alternatives to sodium channel and acetylcholinesterase inhibitors have drawn considerable attention because their new modes of action are more suited to target populations that are currently resistant to insecticides in use in intervention measures ^9–11^.

Two formulations of clothianidin, a neonicotinoid repurposed from the agricultural sector, have been prequalified for indoor residual spraying ^2^. Clothianidin is the unique active ingredient in SumiShield® (Sumitomo Chemical Company, Japan) and is combined with deltamethrin, a pyrethroid, in Fludora Fusion® (Bayer CropScience, Monheim, Germany) ^12– 14^. Neonicotinoids act as agonist of acetylcholine, selectively target the insect nicotinic acetylcholine receptor (nAChR) and disrupt excitatory cholinergic neurotransmission leading to paralysis and death ^15^. Neonicotinoids are intensively used in agriculture and represented more than 25% of the global insecticide sales share in 2014 ^16^. In some Sub-Saharan African countries, between 100 and 200 formulations of thiacloprid, imidacloprid, acetamiprid and thiamethoxam are freely used for agricultural pest management ^17–19^. Neonicotinoids sprayed to protect crops from insect pests are highly water-soluble and are prone to leach in aquatic habitats that support *Anopheles* larvae in farms ^20,21^. This unintentional exposure may contribute to the development of broad-spectrum neonicotinoid resistance ^22–25^. Thus, in prospective areas where clothianidin formulations may be used, it is important to assess baseline susceptibility of vector populations to a wide range of neonicotinoid insecticides. Moreover, bioassays have suggested that other agrochemicals such as thiamethoxam, imidacloprid, and acetamiprid may have satisfactory activity against malaria vectors, but their efficacy on wild populations has yet to be evaluated ^8^.

The present study aimed to test the efficacy of four agrochemicals with or without synergists against malaria mosquitoes. We first evaluated the lethal toxicity of active ingredients and we then compared the susceptibility of wild female adults of two important malaria vector species: *An. coluzzii* and *An. gambia*e. To determine the environmental drivers underlying susceptibility variation within species, we tested several populations from agricultural, rural and urban areas in the equatorial forest of Cameroon. We found that *An. coluzzii* from urban areas is globally susceptible to neonicotinoids while *An. gambiae* is highly tolerant, particularly populations from farming areas. We concluded that neonicotinoid formulations containing adjuvants such as surfactants or other synergists are needed to reach the desired level of efficacy against malaria mosquitoes.

## 2. Materials and methods

### 2.1 Lethal concentrations determination

Center for Disease Control and prevention (CDC) bottle bioassays were used to assess the lethal toxicity of neonicotinoids ^26^. We focused on four active ingredients: acetamiprid, imidacloprid, thiamethoxam and clothianidin. Acetamiprid, imidacloprid and thiamethoxam are commonly used by farmers in Cameroon to protect several types of crops from insect pests ^19,27^. Clothianidin is an agrochemical, which is not registered in Cameroon, but whose formulations have been approved for malaria mosquito control ^2^. All four neonicotinoids tested were technical grade material obtained from Sigma Aldrich Pestanal^®^. These insecticides were dissolved in absolute ethanol except imidacloprid for which acetone was used. A range of concentrations of the active ingredient was tested against one susceptible colony (*An. gambiae* Kisumu or *An. coluzzii* Ngousso) in order to determine LC_50_ and LC_99_ corresponding to the lowest concentrations required to kill respectively 50% and 99% of susceptible mosquitoes. After 1 h exposure to the insecticide, lethal concentrations were determined within 24 h and 72 h. By contrast to clothianidin whose toxicity has been tested with at least one susceptible strain ^7,11,28,29^, there is no information on the lethal concentrations of acetamiprid, imidacloprid and thiamethoxam to African malaria mosquitoes. Therefore, we focused primarily on those three active ingredients. The following gradients were tested for each insecticide: imidacloprid (12.5, 50, 100, 200, 250 µg/ml); acetamiprid (12.5, 25, 50, 75, 150 µg/ml); thiamethoxam (3, 50, 150, 250, 300 µg/ml).

250-ml Wheaton bottles were coated with 1 ml of a given concentration of the insecticide and 25 female adult mosquitoes, 3 to 5 days old, were exposed for 1 h in the bottles. After the exposure period, mosquitoes were removed from the bottles and released into net-covered paper cups on top of which cotton imbibed with 10% sugar solution was placed. Mortality was observed after 24 h and 72 h. Bioassays were performed under a controlled environment of 25–27°C, 70–90% relative humidity and a 12:12 h light/dark photoperiod. Four replicates were tested per concentration together with two controls where mosquitoes were exposed to 1 ml of solvent, ethanol or acetone.

### 2.2 Susceptibility evaluation in wild populations

Wild *An. gambiae sensu lato (s*.*l*.) mosquito populations were collected from several field surveys and cumulatively tested between September 2019 and September 2022. Approval to conduct a study in the Center region (N°: 1-140/L/MINSANTE/SG/RDPH-Ce), ethical clearance (N°: 1-141/CRERSH/2020) and research permit (N°: 000133/MINRESI/B00/C00/C10/C13) were granted by the ministry of public health and the ministry of scientific research and innovation of Cameroon. Mosquitoes were sampled using a dipper from breeding sites identified in eleven locations around Yaoundé. Three locations surveyed within the city were densely urbanized (Tsinga, Combattant, Etoa Meki), while eight sampling sites were located in suburban and rural areas (Figure 1). One of the sites (Nkolondom) was situated in a suburban neighborhood where swampy areas are suitable for intensive cultivation of food crops (Figure 1).

**Figure 1:**
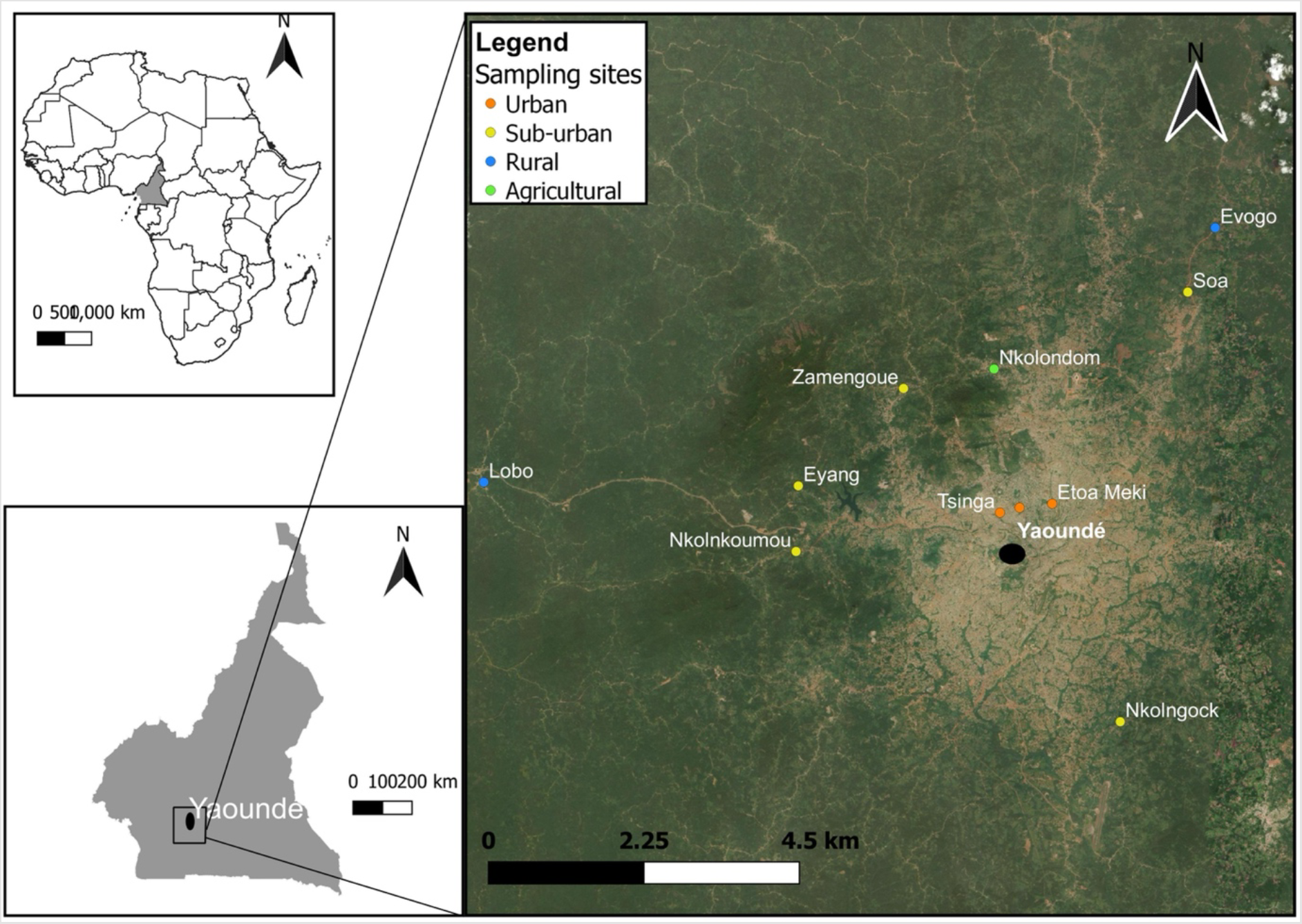
Map of the sampling sites where neonicotinoid susceptibility was monitored in *An. gambiae* and *An. coluzzii*.

Larvae collected from the field were transported in plastic containers to the insectary where they were reared in trays containing 200 ml borehole water. Larvae were given TetraMin® fish food daily, and adults that emerged were maintained in 30 cm-by-30 cm cages and provided with 10% sugar solution. Female adults were tested to assess their level of susceptibility to neonicotinoids using bottle bioassays and a discriminating concentration of the insecticide. A discriminating concentration was defined as the lowest concentration required to kill 100% of susceptible mosquitoes in all four replicates and was chosen within the confidence intervals of LC_99_ for acetamiprid, imidacloprid and thiamethoxam. The discriminating concentration of clothianidin was obtained from ^28^. Two cryptic species of the *An. gambiae* complex: *An. gambiae sensu stricto* (hereafter *An. gambiae*) and *An. coluzzii* are found segregating along an urbanization gradient across the study sites ^30^. The two species were identified from a subset of 50 mosquitoes from each sampling site using a diagnostic PCR method described in ^31^.

### 2.3 Synergistic effect of Piperonyl butoxide (PBO)

A bioassay was performed to determine if the synergistic effect of the cytochrome P450 inhibitor, piperonyl butoxide (PBO) could restore neonicotinoid susceptibility. Mosquitoes that emerged from the same pool of larvae were first exposed to PBO at 4% in CDC bottles, for 1 h before being released into other bottles coated with the neonicotinoid insecticide as described in ^23^. Mortality values obtained with or without prior exposure to the synergist were compared after 72 h of holding period.

### 2.4 Data analysis

Summing the number of dead mosquitoes and expressing this as a percentage of the total number of individuals exposed across all four bottles provided an estimate of the mortality rate per test. Abbott’s formula was used to correct mortality rates when 5%–10% of individuals died in the corresponding control tests ^32^. Average mortality for each insecticide dose and a log-logistic model were used to fit the dose-response curve with the *drc* package in R version 4.2.2 ^33^. A probit model was applied to determine LC_50_ and LC_99_ plus their 95% confidence intervals for each insecticide using the *ecotox* package in R. Mortality in wild populations was interpreted based on the WHO criteria which states that 98%–100% mortality indicates susceptibility, 90%–97% mortality suggests the possibility of resistance that needs to be confirmed and less than 90% mortality corresponds to resistance ^34^.

## 3. Results

### 3.1 Lethal toxicity of neonicotinoids to *Anopheles*

The short-term lethal toxicity of three neonicotinoids against *Anopheles* mosquitoes was tested with dose-response bioassays using four replicates per dose of insecticide. Results of LC_50_ and LC_99_ of acetamiprid, thiamethoxam and imidacloprid are presented in Table 1 and Figure 2 showing the dose-response curves with mortality recorded within 24 h and 72 h of exposure. 24-h LC_99_ obtained with the susceptible strain *An. gambiae* Kisumu were 69.0 µg/ml, confidence interval (*CI*)_95%_ [54.40, 98.1], for acetamiprid compared to 152.0 µg/ml [112.0, 235.1] for imidacloprid. Meanwhile, 24-h lethal concentrations of thiamethoxam determined with the susceptible strain *An. coluzzii* Ngousso were LC_50_: 9.6 µg/ml [6.8, 13.5] and LC_99_: 133.0 µg/ml [79.4, 277.0], respectively. When toxicity is high, lethal concentrations are low since smaller doses of insecticide are required to kill a given number of exposed individuals. Based on the lowest LC_50_, thiamethoxam (LC_50_: 9.6 µg/ml [6.8, 13.5]) was the most toxic neonicotinoid to *Anopheles* mosquitoes in 24 h followed by acetamiprid (LC_50_: 13.6 µg/ml [11.5, 15.5]) and imidacloprid (LC_50_: 18.6 µg/ml [15.2, 20.0]). To compare lethal toxicity within 24 h and 72 h, we used the 95% confidence intervals. The mortality response of populations was considered different between 24 h and 72 h if there was no overlap between their corresponding 95% confidence limits. Confidence intervals of LC_50_ and LC_99_ at 24 h and 72 h overlapped for the three active ingredients tested, which suggested that extending the holding period did not increase toxicity (Table 1).

**Table 1:** Lethal concentrations, LC_50_ and LC_99_, estimated at 24 h and 72 h post-exposure

| Active ingredient | Species | Number of mosquitoes | Holding period | LC<sub>50</sub> value (µg/L) | Lower-upper 95% confidence intervals (µg/L) | Standard error | LC<sub>99</sub> value (µg/L) | Lower-upper 95% confidence intervals (µg/L) | Standard error | Chi Square |
| --- | --- | --- | --- | --- | --- | --- | --- | --- | --- | --- |
| acetamiprid | *An. gambiae* (Kisumu) | 500 | 24 h | 13.6 | 11.5 - 15.5 | 1.08 | 69 | 54.4 - 98.1 | 1.16 | 18.1 |
|  |  |  | 72 h | 13.9 | 11.3 - 16.2 | 1.07 | 64.1 | 48.7 - 101 | 1.15 | 24.3 |
| imidacloprid | *An. gambiae* (Kisumu) | 350 | 24 h | 18.6 | 15.2 - 22.0 | 1.1 | 152 | 112 - 235 | 1.2 | 18.4 |
|  |  |  | 72 h | 17.1 | 12.9 - 21.4 | 1.1 | 148 | 100 - 273 | 1.21 | 22 |
| thiamethoxam | *An. coluzzii* (Ngouso) | 475 | 24 h | 9.66 | 6.78 - 13.5 | 1.14 | 133 | 79.4 - 277 | 1.26 | 25.7 |
|  |  |  | 72 h | 7.47 | 5.84 - 9.67 | 1.13 | 75.3 | 47.8 - 142 | 1.31 | 13.2 |

**Figure 2:**
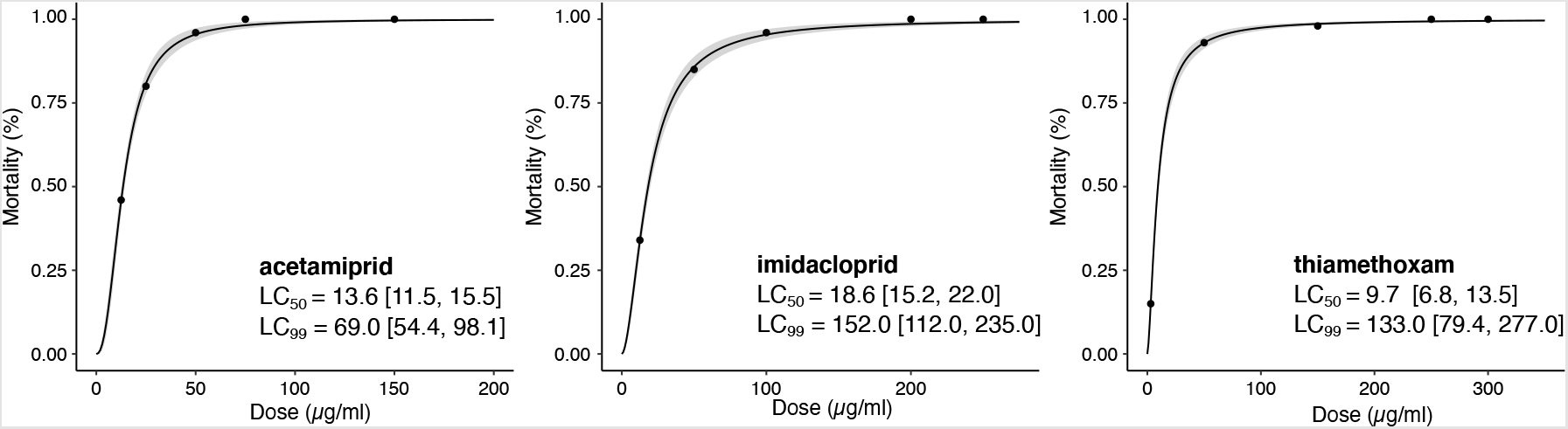
Dose-response curves with the standard error of the regression model (grey bands) representing the 24-h toxicity of three neonicotinoids in *Anopheles* mosquitoes. LC_50_ and LC_99_ with 95% confidence intervals of acetamiprid and imidacloprid were determined using the susceptible strain *An. gambiae* Kisumu while lethal concentrations of thiamethoxam were estimated against the *An. coluzzii* Ngousso strain.

### 3.2 Variation in susceptibility within and between species

To evaluate the baseline susceptibility of wild populations to acetamiprid, imidacloprid, and thiamethoxam, we chose discriminating concentrations of 75 µg/ml, 200 µg/ml and 250 µg/ml, respectively. The doses were picked up within the 95% confidence intervals for 24-h LC_99_ in order to balance the risk of not detecting low-level resistance while limiting the risk of reporting false positives. The doses of acetamiprid and imidacloprid we chose were recently used to assess the susceptibility of wild *An. coluzzii* populations and have revealed satisfactory discriminative power ^22^. In addition to acetamiprid, imidacloprid and thiamethoxam, we tested a fourth neonicotinoid (clothianidin) using a discriminating concentration of 150 µg/ml as determined in an earlier study ^28^. We first confirmed the efficacy of the discriminating concentrations against the insecticide susceptible strains *An. gambiae* Kisumu and *An. coluzzii* Ngousso. For both laboratory colonies, 100% of female adults exposed to the discriminating dose of each of the four neonicotinoids died within 24 h. The baseline susceptibility of the two major vectors *An. gambiae and An. coluzzii* to acetamiprid, imidacloprid and thiamethoxam was determined from populations tested from 2020 to 2022. Susceptibility to clothianidin was estimated by pooling original data obtained from bioassays carried out between 2020 and 2022 with data presented in ^23^, which originated from field collections conducted from 2019 to 2020. In total, 5532 female mosquitoes belonging to eleven populations were tested with bioassays from 2019 to 2022.

We first analyzed bioassay results from individual collection sites. We observed that populations collected from three sites situated in urbanized areas were 100% *An. coluzzii* and were susceptible to neonicotinoids except acetamiprid for which signs of reduced susceptibility were apparent (Figure 3). 100% mortality was observed within 24 h or 72 h holding period upon exposure to thiamethoxam, imidacloprid and clothianidin. Meanwhile, average 72-h mortality to acetamiprid was 80% in *An. coluzzii* collected from the urban neighborhoods. Across two sites located in the suburban area (Nkolkoumou and Soa), the relative frequencies of *An. gambiae* and *An. coluzzii* were ∼80% and 20%, respectively. Nkolnkoumou mosquito populations were susceptible to imidacloprid and clothianidin and resistant to thiamethoxam and acetamiprid. Soa samples were tested only with clothianidin and were susceptible. None of the populations from typical *An. gambiae* habitats (rural areas) was susceptible to acetamiprid or clothianidin. Populations from the agricultural site Nkolondom were the least susceptible to neonicotinoids, with mortality rates below 50% for acetamiprid and clothianidin. It was also the only site where resistance to all four neonicotinoids was detected.

**Figure 3:**
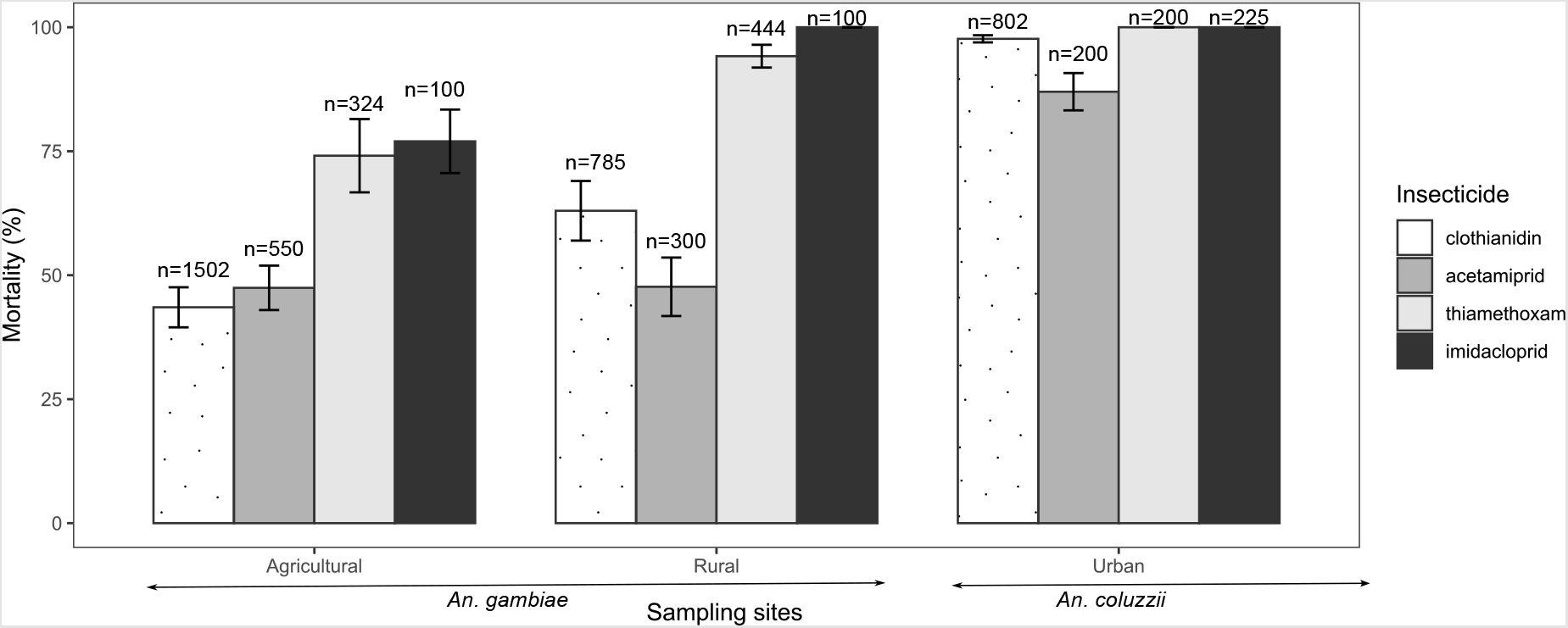
Baseline susceptibility of 5532 female adults of *An. gambiae* and *An. coluzzii* to four neonicotinoids obtained from pooled data representing 4 years of monitoring. Error bars indicate the standard error of the mean and (n) the number of mosquitoes tested.

We then pooled results from all the bioassay tests carried out over a four-year monitoring period and we compared susceptibility profiles between urban, rural and agricultural areas. It appeared that *An. gambiae* population collected from farms were the most tolerant and that agricultural pest management is likely spreading neonicotinoid resistance among neighboring rural populations (Figure 3). Results from the two suburban sites harboring 80% *An. gambiae* were pooled with those of six locations that were typically rural and contained 100% *An. gambiae*. While thiamethoxam was the most toxic to *Anopheles* mosquitoes based on the lowest LC_50_, imidacloprid was the most effective considering mortality induced in wild populations. Clothianidin, which is neither an agricultural nor a public health insecticide in Cameroon induced low mortality in *An. gambiae* populations from rural and agricultural areas, which suggests that locally used neonicotinoids or other mechanisms are conferring cross-resistance to clothianidin.

### 3.3 Synergistic effect of piperonyl butoxide (PBO)

Adjuvants such as vegetable oil surfactants drastically enhance the potency of neonicotinoids against *Anopheles* mosquitoes ^25^. Bioassays were carried out to determine if inhibition of cytochromes mediated by PBO could also improve the efficacy of some active ingredients. We tested the synergistic effect of PBO using the agricultural population that was resistant to the four neonicotinoids. We also focused on clothianidin, acetamiprid and thiamethoxam because mortality rates within 72 h of exposure were less than 75% allowing a more accurate evaluation of synergism ^34^. Susceptibility was fully restored for acetamiprid (100% vs 35.55 ± 5.25%) and partially restored for clothianidin in the presence of PBO (74.33 ± 3.82 vs 30.98 ± 3.49, Wilcoxon rank sum test, p<0.05) after 72 h post-exposure (Figure 4). On the contrary, pre-exposure to PBO did not significantly affect susceptibility to thiamethoxam (58.0% ± 8.2 vs 71.5% ± 7.7, Wilcoxon rank sum test, p>0.05).

**Figure 4:**
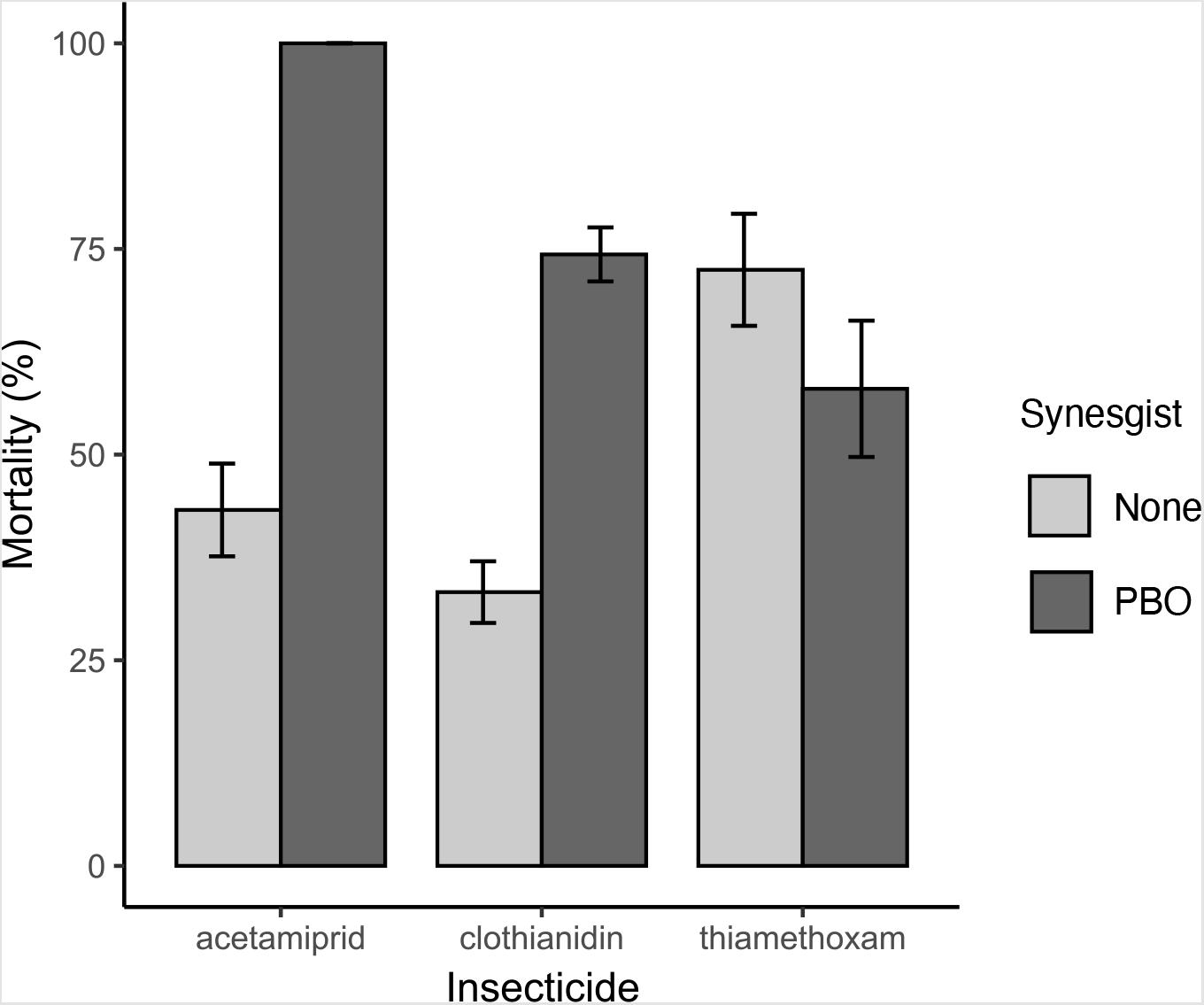
Effects of piperonyl butoxide (PBO) on the efficacy of three neonicotinoids against resistant *An. gambiae* mosquitoes.

## Discussion

Most insecticides used in mosquito control come from the agricultural sector ^4^. A new generation of active ingredients are being tested to control anopheline mosquitoes that have become highly resistant to existing vector control insecticides ^7,8^.

In the present study, we focused on neonicotinoids, a class of insecticides including clothianidin whose formulations are proposed for indoor residual spraying ^12–14^. We first determined the lethal concentrations of three different neonicotinoids using two susceptible laboratory colonies. 24-h lethal concentration (LC_99_) indicated that acetamiprid is the most toxic neonicotinoid to anopheline mosquitoes as it had the lowest LC_99_, followed by clothianidin ^28^. However, in comparison with some public health insecticides, the neonicotinoids we tested had substantially lower toxicity based on LC_99_ values. For example, 24-h LC_99_ of the pyrethroid, deltamethrin, to *Anopheles* is approximately 7-fold lower than that of the most potent neonicotinoid, acetamiprid ^26^. A large-scale screening conducted to search for candidate pesticides that could be used for malaria vector control revealed that three neonicotinoids were among the least active insecticides against adults of *Aedes aegypti* and *Anopheles stephensi* in a list of nearly 100 compounds tested ^8^. LC_80_ of > 200 µg/ml was obtained in 24 h when an insecticide-susceptible strain of *An. stephensi* was exposed to imidacloprid or thiamethoxam while LC_80_ for acetamiprid was ∼20 µg/ml ^8^. In the current study, we also noted that extending the holding period post-exposure did not improve the toxicity of neonicotinoids as there was no significant difference between LC_50_ and LC_99_ at 24 h and 72 h.

In addition to relatively low toxicity, there is also ongoing reduction in susceptibility in wild mosquito populations. Based on a 4-year monitoring and testing of adult populations from 11 sites, we found contrasting patterns of baseline susceptibility to neonicotinoids between the two sibling species *An. gambiae* and *An. coluzzii*. Here we applied discriminating doses that were determined using two insecticide susceptible strains. The doses have shown sufficient discriminative power and were able to reveal gradients of tolerance to neonicotinoids in wild populations of two important vector species.

Our study corroborated findings from earlier surveys indicating that clothianidin resistance is emerging in *An. gambiae* despite the fact this active ingredient remains an exotic insecticide in Cameroon ^23,24^. By contrast, it has been reported that formulations of acetamiprid and imidacloprid are intensively used for crop protection in agricultural areas in Cameroon ^19,23,27^. As neonicotinoids are highly soluble in water, they are likely leaching in aquatic habitats in farms and are contributing to selecting for resistance in anopheline populations ^20,21^. In line with this prediction, a study revealed that *An. gambiae* larvae collected from Nkolondom retained high fitness when reared in water containing concentrations of neonicotinoids that were lethal to susceptible strains ^24^ The present study shows that neonicotinoid tolerance selected at larval stage induces cross-resistance to several active ingredients in female adult mosquitoes that are vectors of *Plasmodium* ^23–25^. *An. gambiae*, which occurs in the countryside where agricultural activities are more frequent and are associated with intensive use of pesticides are developing resistance to several neonicotinoids ^19,27^. Precisely, we noted that resistance was strongest in samples collected from the agricultural areas, where formulations of neonicotinoids were being used. *An. coluzzii* populations collected from urbanized areas of Yaoundé where neonicotinoids are less or not used had sub-optimal mortality (∼80%) to acetamiprid, but this species was generally susceptible to neonicotinoids. This finding was once again consistent with larval tests, which showed that third instar larvae of *An. coluzzii* from Yaoundé had low survival and barely emerged in water contaminated with neonicotinoids ^24^. However, in Ivory Coast, reduced susceptibility to acetamiprid and imidacloprid has been reported in adults of *An. coluzzii* collected from agricultural areas suggesting that populations of this species can develop resistance if they are exposed chronically to neonicotinoid residues ^22^.

Our study highlights the need for a more comprehensive survey of susceptibility to neonicotinoids in malaria vectors species, especially in prospective areas where clothianidin formulations may be used ^35^. The vectorial system in Sub-Saharan Africa is complex and comprises at least a dozen major species that occupy diverse niches at larval and adult stages ^36,37^. Our results show that, depending on the history of exposure to agricultural neonicotinoids, the susceptibility profile could vary significantly within and between *Anopheles* species, even on a small geographic scale. The species whose larvae are more likely to be exposed to neonicotinoids in their native ranges are *An. gambiae, An. coluzzii* and *An. arabiensis* as they are known to exploit man-made habitats such as temporary breeding sites created in farms ^38,39^. These species should be the focus of extensive monitoring and efforts. *An. funestus* is a very important vector whose populations should be carefully scrutinized even if larval habitats of this species are less prone to anthropogenic disturbance ^40^.

Surfactants including cleaning products such as soap have potential to synergize neonicotinoid efficacy in malaria vectors ^7,22,25,41^. This synergism provides an opportunity to improve neonicotinoid formulations that could be used vector control. In the present study, we have revealed that PBO is another effective synergist, which offers additional options to enhance neonicotinoid efficacy against *Anopheles*. We have observed that when resistant *Anopheles* mosquitoes collected from farms were pre-exposed to PBO, they became susceptible to acetamiprid suggesting that cytochrome P450 monooxygenases are involved in resistance to this insecticide. Indeed, overexpression of cytochrome P450s monooxygenases is an important mechanism underlying neonicotinoid resistance in insect pests including mosquitoes, white fly and aphids ^23,42,43^. Gene expression analysis revealed overexpression of multiple P450 genes in an acetamiprid-resistant strain of the melon aphid, *Aphis gossypii*, indicating a role of P450-mediated detoxification in acetamiprid resistance ^44^. Acetamiprid resistance has also been shown to depend strongly on monooxygenases in white flies and *Aedes* mosquitoes ^45,46^. Consistent with past surveys, pre-exposure to PBO also drastically improve the efficacy of clothianidin against *Anopheles* mosquitoes ^23^. One notable exception was thiamethoxam as PBO had little effect on baseline susceptibility of wild populations. This could be explained by the fact that thiamethoxam is a pro-insecticide which needs to be converted to clothianidin before being active in insects. This conversion is catalyzed by enzymes which might have been inhibited by PBO, leading to a slight reduction in insecticidal activity when mosquitoes were pre-exposed to PBO ^47^.

In conclusion, although neonicotinoids have low acute toxicity and reduced efficacy in some *Anopheles* mosquito populations, potent formulations can still provide alternatives to control pyrethroid-resistant malaria mosquitoes. In the current study and complimentary investigations ^25^, we have found that available synergists such as soap or PBO could be used to enhance the efficacy of neonicotinoid insecticides against *Anopheles* mosquitoes.

## Author Contributions

FA: Conceptualization, Formal analysis, Investigation, Methodology, Writing – original draft; CF: Conceptualization, Formal analysis, Investigation, Methodology; MA: Investigation, Methodology; VP-B, Resources, Supervision. CK: Conceptualization, Formal analysis, Funding acquisition, Investigation, Project administration, Writing – review & editing.

## Informed Consent Statement

Not applicable.

## Data Availability Statement

The data for this study have been presented within this article.

